# An ancient lineage of highly divergent parvoviruses infects both vertebrate and invertebrate hosts

**DOI:** 10.1101/571109

**Authors:** Judit J Pénzes, William Marciel de Souza, Mavis Agbandje-McKenna, Robert J. Gifford

**Affiliations:** University of Florida, McKnight Brain Institute and Department of Biochemistry and Molecular Biology, 1149 Newell Dr, Gainesville, FL 32610, USA; Virology Research Center, School of Medicine of Ribeirão Preto of the University of São Paulo, Ribeirão Preto, Brazil; MRC-University of Glasgow Centre for Virus Research, 464 Bearsden Road, Glasgow, UK

## Abstract

Chapparvoviruses are a highly divergent group of parvoviruses (family *Parvoviridae*) first identified in 2013. Interest in these poorly characterized viruses has been raised by recent studies indicating that they are the cause of chronic kidney disease that arises spontaneously in laboratory mice. In this study, we investigate the biological and evolutionary characteristics of chapparvoviruses via comparative analysis of genome sequence data. Our analysis, which incorporates sequences derived from endogenous viral elements (EVEs) as well as exogenous viruses, reveals that chapparvoviruses are an ancient lineage within the family *Parvoviridae*, clustering separately from members of both currently established parvoviral subfamilies. Consistent with this, they exhibit a number of characteristic genomic and structural features, i.e. a large number of putative auxiliary protein-encoding genes, capsid protein genes non-homologous to any hitherto parvoviral *cap*, as well as a putative capsid structure lacking the canonical fifth strand of the ABIDG sheet comprising the luminal side of the jelly roll. Our findings demonstrate that the chapparvovirus lineage infects an exceptionally broad range of host species, including both vertebrates and invertebrates. Furthermore, we observe that chapparvoviruses found in fish are more closely related to those from invertebrates than they are to those that infect amniote vertebrates. This suggests that transmission between distantly related host species may have occurred in the past. Our study provides the first integrated overview of the chapparvovirus group, and revises current views of parvovirus evolution

**AUTHOR SUMMARY:** Chapparvoviruses are a recently identified group of viruses about which relatively little is known. However, recent studies have shown that these viruses cause disease in laboratory mice and are prevalent in the fecal virome of pigs and poultry, raising interest in their potential impact as pathogens, and utility as experimental tools. We examined the genomes of chapparvoviruses and endogenous viral elements (‘fossilized’ virus sequences derived from ancestral viruses) using a variety of bioinformatics-based approaches. We show that the chapparvoviruses have an ancient origin and are evolutionarily distinct from all other related viruses. Accordingly, their genomes and virions exhibit a range of distinct characteristic features. We examine the distribution of these features in the light of chapparvovirus evolutionary history (which we can also infer from genomic data), revealing new insights into chapparvovirus biology.

## INTRODUCTION

Parvoviruses (family *Parvoviridae*) are small, non-enveloped viruses with T=1 icosahedral symmetry and linear, single-stranded DNA (ssDNA) genomes ∼4-6 kilobases (kb) in length. The family has historically been divided into two subfamilies; *Parvovirinae* and *Densovirinae*, containing viruses that infect vertebrate and invertebrate hosts, respectively (1). Despite exhibiting great variation in expression and transcription strategies, they have a relatively conserved overall genome structure: a non-structural (NS) expression cassette is located at the left side of the genome, while the structural viral proteins (VPs) are encoded by the right, and complex, hairpin-like DNA secondary structures are present at both genomic termini (2). Small satellite proteins and an assembly-activating protein have been discovered as products of open reading frames overlapping the right hand expression cassette (3, 4)

Numerous novel parvoviruses have been identified in recent years, primarily via approaches based on high throughput sequencing (HTS) (5-11). In addition, progress in whole genome sequencing of eukaryotes has revealed that sequences derived from parvoviruses occur relatively frequently in animal genomes (12-17). These endogenous parvoviral elements (EPVs) are derived from the genomes of ancient parvoviruses that were incorporated into the gene pool of ancestral host species. This can presumably occur when infection of a germline cell leads to parvovirus-derived DNA becoming integrated into host chromosomes, and the cell containing the integrated sequences then goes on to develop into a viable organism (18). Many EPVs are millions of years old, and are genetically “fixed” in the genomes of host species (i.e. all members of the species have the integrated EPV in their genomes). Such ancient EPV sequences are in some ways analogous to “parvovirus fossils”, since they preserve information about the ancient parvoviruses that infected ancestral animals.

Among the novel parvovirus groups identified via sequencing, one - provisionally labeled ‘chapparvovirus’ - stands out as being particularly unusual. These viruses, which have been primarily reported via metagenomic sequencing of animal feces, derive their name from an acronym (CHAP), referring to the host groups in which they were first identified (<u>CH</u>iropteran- <u>A</u>vian-<u>P</u>orcine) (15, 16, 19, 20). Subsequently several additional chapparvovirus sequences have been reported, including some that were identified in whole genome sequence (WGS) data derived from vertebrates, including reptiles, mammals, and birds (9). These sequences were picked up by *in silico* screens designed to detect EPV. However, since all the chapparvovirus sequences identified in WGS data lack clear evidence of genomic integration, it is thought likely that they actually derive from infectious chapparvovirus genomes that contaminated WGS samples, rather than from EPVs (9).

Until relatively recently, evidence that the chapparvoviruses detected via sequencing actually infect vertebrate hosts has been lacking. However, a recent study has claimed to demonstrate that a chapparvovirus called ‘mouse kidney parvovirus’ (MKPV) circulates among laboratory mice populations, in which it causes a kidney disease known as ‘inclusion body nephropathy’ (21). These findings imply that chapparvoviruses represent a potential disease threat to humans and domestic species. In addition, they have raised interest in the use of these viruses as experimental tools.

In this study, we perform comparative analysis of ChPV genomes and ChPV-derived EPVs, revealing new insights into the biology and evolution of this poorly understood group.

## RESULTS

### Identification and characterization of novel chapparvovirus sequences

We systematically screened published WGS data and identified a total of fifteen previously unreported, chapparvovirus-derived DNA sequences. Two were identified in vertebrates and thirteen in invertebrates (**Table 1**). The majority of the novel chapparvorvirus sequences identified in our screen were derived from the replicase (*rep*) gene, but we identified complete sequences derived from both the *rep* and capsid (*cap*) genes in two species: the Gulf pipefish (*Syngnathus scovelli*) and the black widow spider (*Latrodectus hesperus*). Partial *cap* genes were identified in the scarab beetle (*Oryctes borbonicus*), taurus scarab (*Onthophagus taurus*) and Chinese golden scorpion (*Mesobuthus martensii*) elements (**Figure 1**).

**Table 1.**
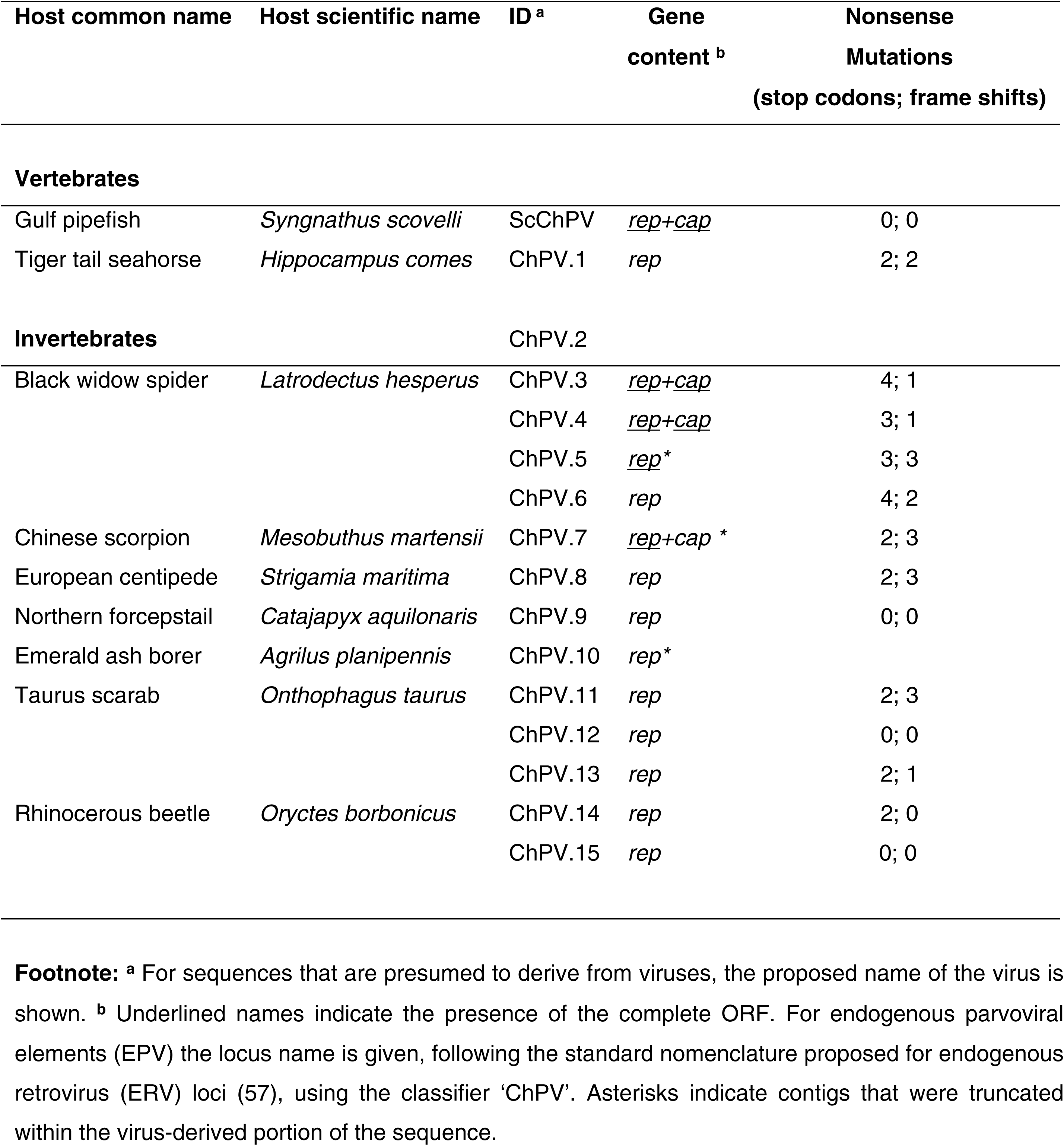
Novel chapparvovirus sequences identified in this study.

**Figure 1.**
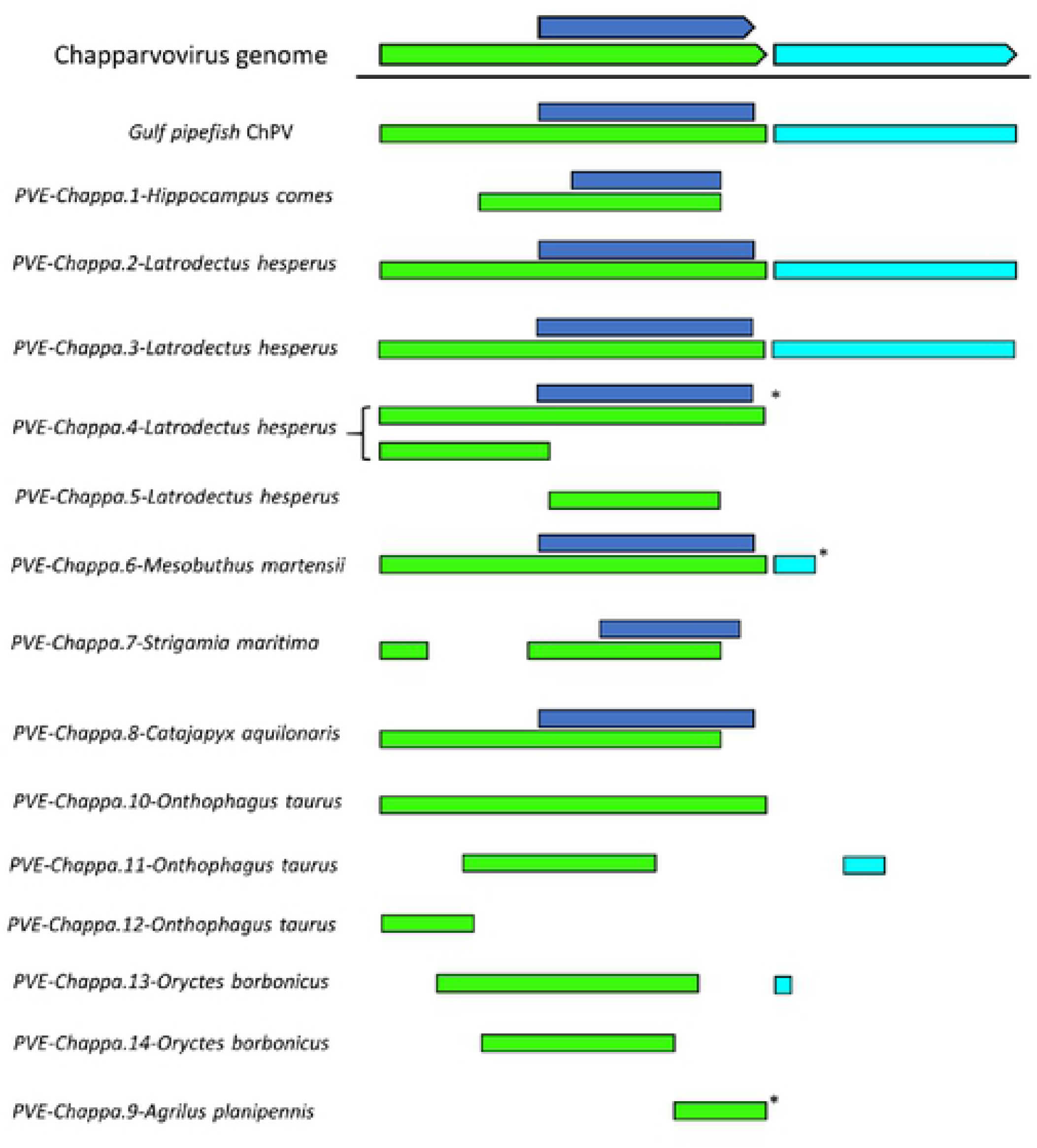
Basic gene content of novel chapparvoviruses and chapparvovirus EPV, shown in relation to a representative chapparvovirus genome (mouse kidney parvovirus). Asterisks indicate contigs that were truncated within the virus-derived portion of the sequence. Abbreviations: non-structural protein (NS); capsid protein (VP); nucleoprotein (NP).

Two of the novel sequences were identified in WGS assemblies of fish in the family Syngnathidae, including the tiger tail seahorse (*Hippocampus comes*) and the Gulf pipefish. The seahorse sequence clearly represented an EPV (**Figure 2a**), while the pipefish sequence lacked flanking genomic sequences and appeared to derive from an exogenous virus. In addition, all hits of invertebrate origin clearly represented EPVs (**Figure 2a**). However, none shared homologous flanking sequences, suggesting that they each derive from distinct germline incorporation events.

**Fig.2 legend:**
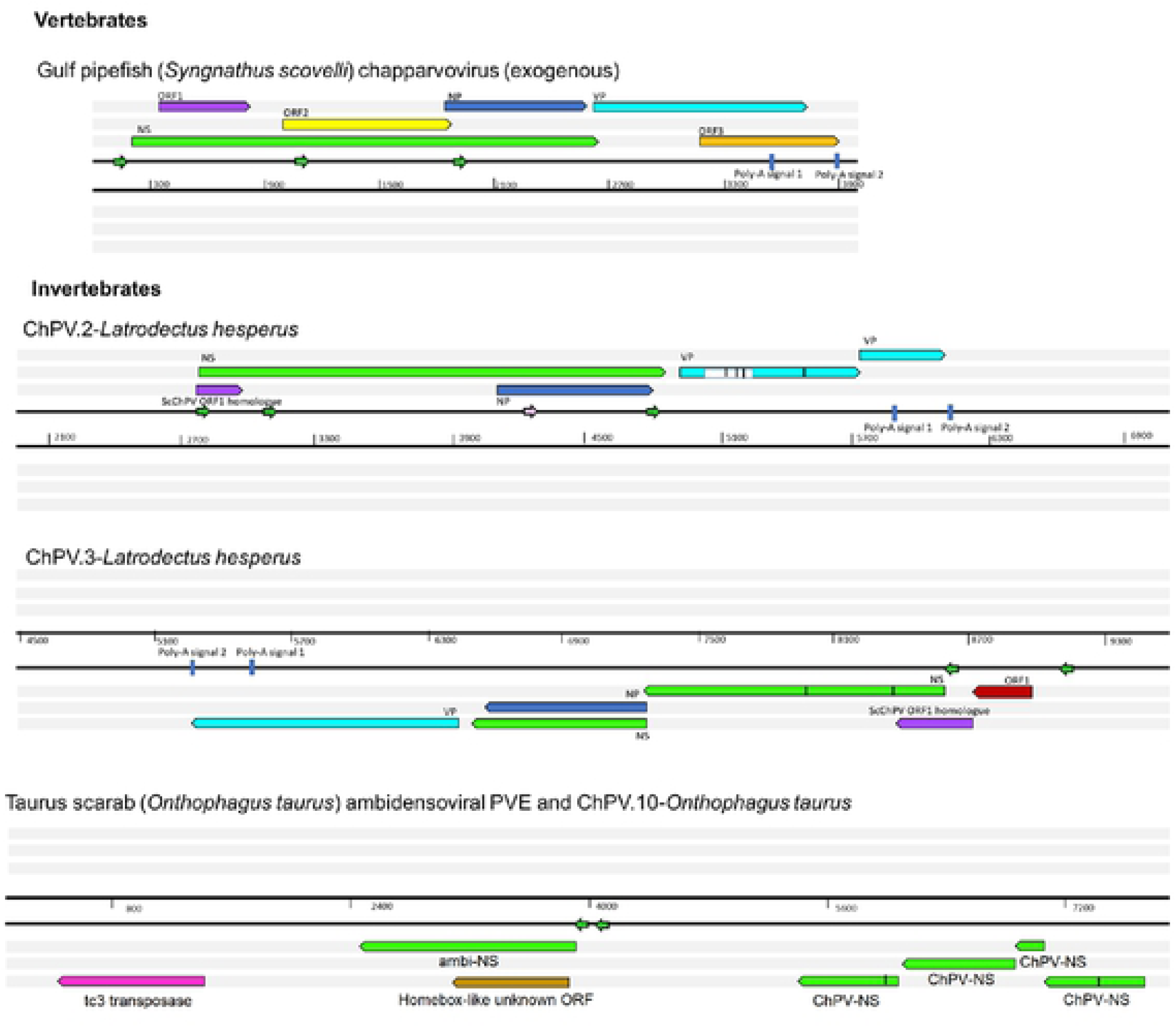
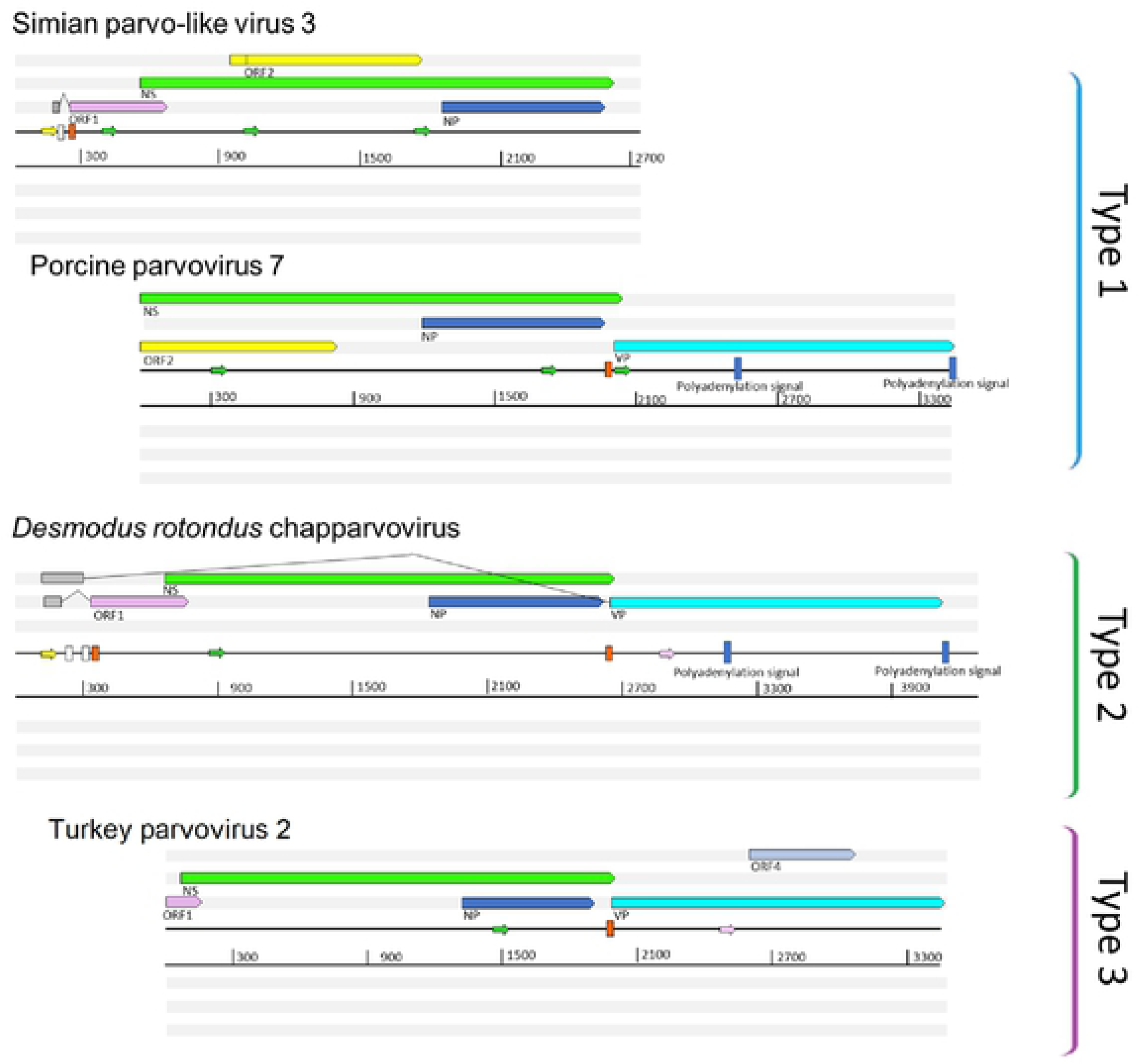
Genome organization, position of open reading frames (ORFs) and predicted cis transcription elements of chapparvoviruses and endogenous chapparvoviral elements shown in six reading frames. ORFs are represented by arrows, colored according to homology. In-frame stop codons are shown as vertical lines. Splice donor sites are marked by white-colored, acceptor sites by orange-colored bars. Blue colored bars show predicted polyadenylation signals. Promoters are presented as small green arrows if predicted with a higher score than 0.95 of 1 and pink if the score is between 0.9 and 0.95. Sequences marked by grey boxes are assessed to be transcribed but not translated. (A) Near complete entries detected include the exogenous, potentially circulating chapparvovirus of the gulf pipefish (*Syngnathus scovelli*) as well as two, non-identical, endogenous chaparvoviruses of the Western black widow spider (*Latrodectus hesperus*). ChPV.2-Latrodectus hesperus contains a previously unidentified repetitive element, present as multiple copies scattered in the *Latrodectus* genome, marked by the white box within the VP gene. The element ChPV.10-Onthophagus taurus shares its integration site with another endogenous element disclosing similarity to ambidensoviruses. (B) Representatives of the three basic genome organization types of exogenous amniote chapparvoviruses. ORF4, however, does not occur in any of the other avian chapparvoviruses.

We used phylogenetic approaches to reconstruct the evolutionary relationships of newly identified chapparvovirus sequences to previously reported parvoviruses. We reconstructed evolutionary relationships using maximum likelihood-based approaches and an alignment spanning the tripartite helicase domain of the NS protein (**Figure 3a**). Strikingly, reconstructions indicated that the family *Parvoviridae* four major clades, rather than the two that have historically been recognised (1). Of the four major lineages, one corresponds to the subfamily *Parvovirinae* as recognised by current taxonomic schemes. However, the subfamily *Densovirinae* appeared to be split into two clades; one encompassing all ambisense densoviruses along with viruses of the genus *Iteradensovirus* (which have monosense genomes) and the second comprised by genera *Hepan*-, *Brevi*- and *Penstyldensovirus*. Moreover, a fourth parvovirus lineage was evident, comprised of the ChPVs and ChPV-derived EPVs.

**Figure 3 legend:**
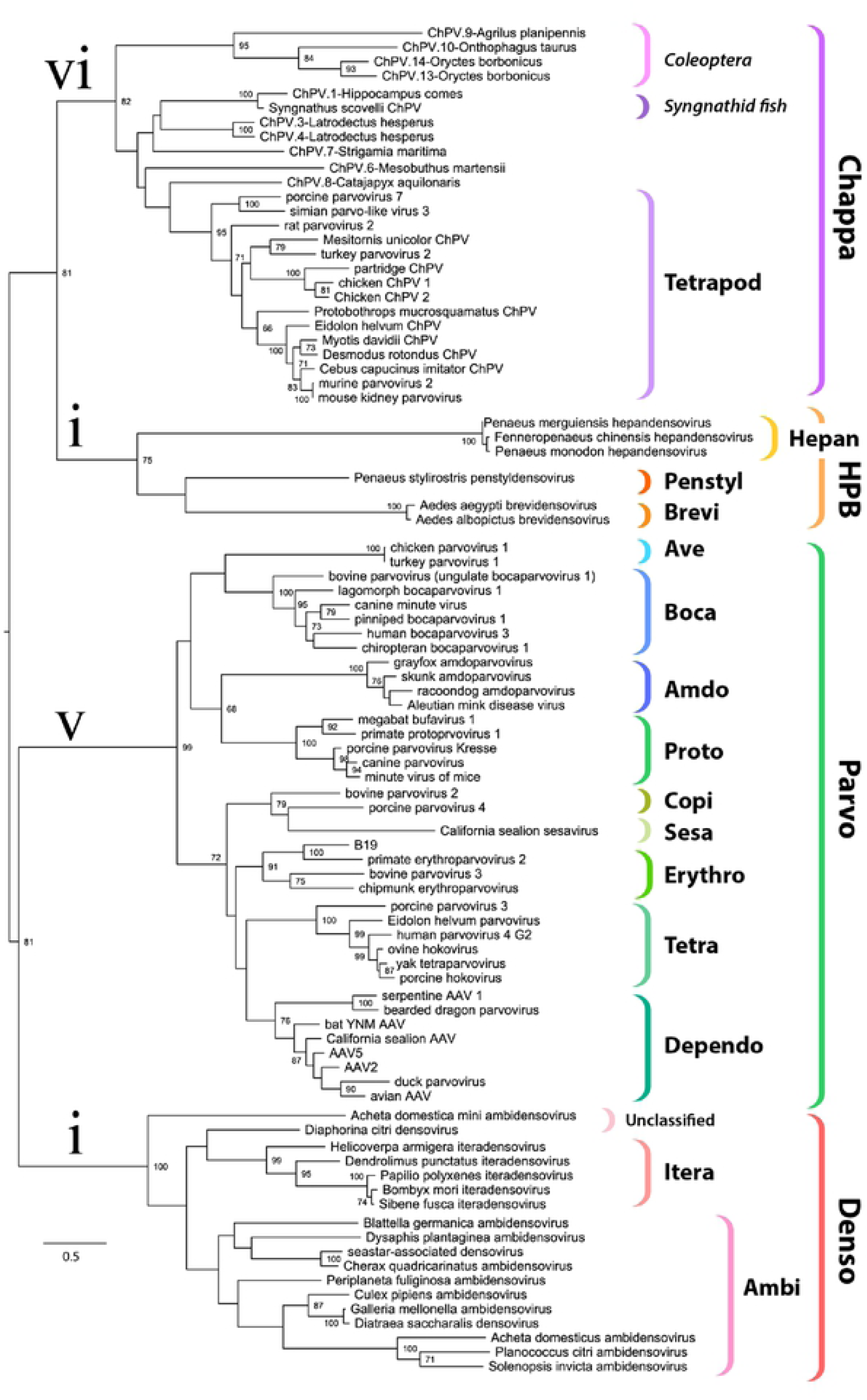
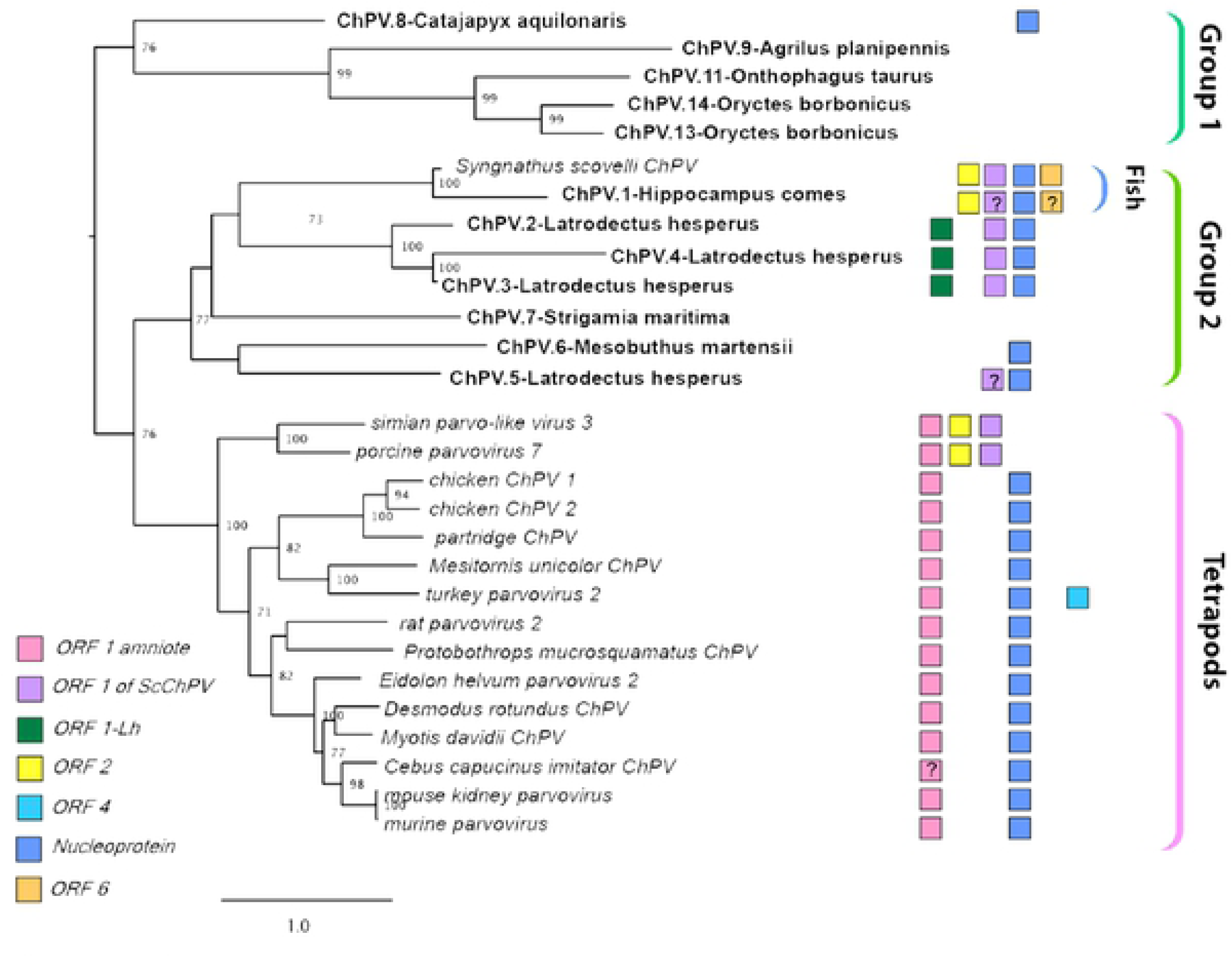
(A) Maximum likelihood phylogeny of the family *Parvoviridae* based on the tripartite helicase domain of the large non-structural protein NS1. (B) Maximum likelihood phylogenetic reconstructions of the Chapparvovirus clade based on the complete aligned amino acid sequences of the NS1. The presence or suspected presence (marked by a question mark) of predicted auxiliary protein encoding open reading frames (ORFs) are mapped as boxes of various colors next to each entry. Bold taxa labels indicate endogenous sequences, whereas taxa labels in italics indicate sequences known or believed to derive from viruses.

The branching relationships among ChPVs were not fully resolved by phylogenetic analysis of the helicase domain. The putative large replicase proteins (NS1) of ChPVs displayed a high degree of amino acid variability, particularly toward their N- and C-termini. However, a region ∼500-aa-long could be aligned reliably throughout all complete and partial entries previously proven to cluster within the chapparvovirus lineage in case of the NS helicase-based inference. According to this, the ChPV clade is comprised of three significantly-supported monophyletic lineages. One of these includes ChPVs sampled from amniotes (reptiles, birds, and mammals), in which three further consistently well-supported sublineages could be observed (**Figure 3b**). The amniote ChPVs form a sister clade to EPVs found in the arthropod subphyla Chelicerata (arachnids, camel spiders, scorpions, whip scorpions, harvestmen, horseshoe crabs and kin) and Myriapoda (millipedes, centipedes and kin) as well as with the syngnathe fish associated sequences. Within this clade, however, only the grouping of spider and syngnathe fish sequences was supported. A third lineage was also observed, containing sequences from the arthropod subphylum Hexapoda (insects, springtail, and forcepstail). Monophyly of the beetle EPVs is robustly supported (**Figure 3b**).

### Comparative analysis of previously reported chapparvovirus genomes

We performed comparative analysis of nine previously sequenced ChPV genomes so that we might: (i) identify genome features that characterise these viruses, and (ii) make inferences about aspects of ChPV biology and evolution (**Figure 2b**). ChPV genomes tend toward the shorter end of the parvovirus genome size range (∼4kb). They encode a relatively long *rep* gene, and a relatively short *cap* gene. The *rep* gene product (NS) is ∼650 amino acids (aa) in length, with the longest example being the 668 aa long protein encoded by *Desmodus rotondus* chapparvovirus (DrChPV). Chapparvovirus NS proteins contain ATPase and helicase domains, but these are the only regions exhibiting clear homology to those found in other parvovirus groups (**Figure 2b**). Overlapping the *rep*, a predicted minor ORF, ∼220 aa in length, is located in a position equivalent to that of the nucleoprotein (NP) ORF found in certain *Parvovirinae* genera (i.e. *Ave*- and *Bocaparvovirus*). However, it should be noted that the polypeptide encoded by this gene – which we tentatively refer to as NP - exhibits no significant similarity to any other parvovirus NP proteins. Secondary structure predictions indicate that the vast majority of the NP protein has a mostly helical structure, with numerous possible phosphorylation sites as well as a potentially protein-binding disordered N-term (**Figure S2**). Together, these observations suggest a non-structural function. The NP ORF, although of similar length in all genomes, has no canonical start codon in case of porcine parvovirus 7 (PPV7) and simian parvo-like virus 3 (SPV3). This would imply that in these viruses, splicing of the *rep* RNA is required for expression of the NP protein.

All chapparvoviruses appear to be characterized by relatively short VP sequences that are between 450- 500 aa in length, as opposed to ∼650-820 aa as found in most other parvoviruses (the exception being the brevi- and penstyldensoviruses which encode an even shorter VP). In all chapparvoviruses, the first methionine of the VP ORF is preceded by potential coding sequence, and in all published ChPV sequences, a canonical splice acceptor site is located immediately upstream. Possibly, the VP ORF encodes only the major capsid protein, and there may be other versions of this VP protein that are elongated at the N-terminus, and are incorporated into the capsid at a lower copy number, as found in the majority of parvoviruses (1). However, the only splice donor sites we identified are located relatively far upstream. In MKDV, however, there are two large introns present, putting these upstream exons in frame with the VP encoding exon.

The VP proteins of chapparvoviruses share no significant sequence similarity with those of other parvoviruses. Interestingly, however, structural similarity with erythro-, proto, and bocaparvoviruses can be detected for VP using fold recognition (**Figure 4**).

**Figure 4.**
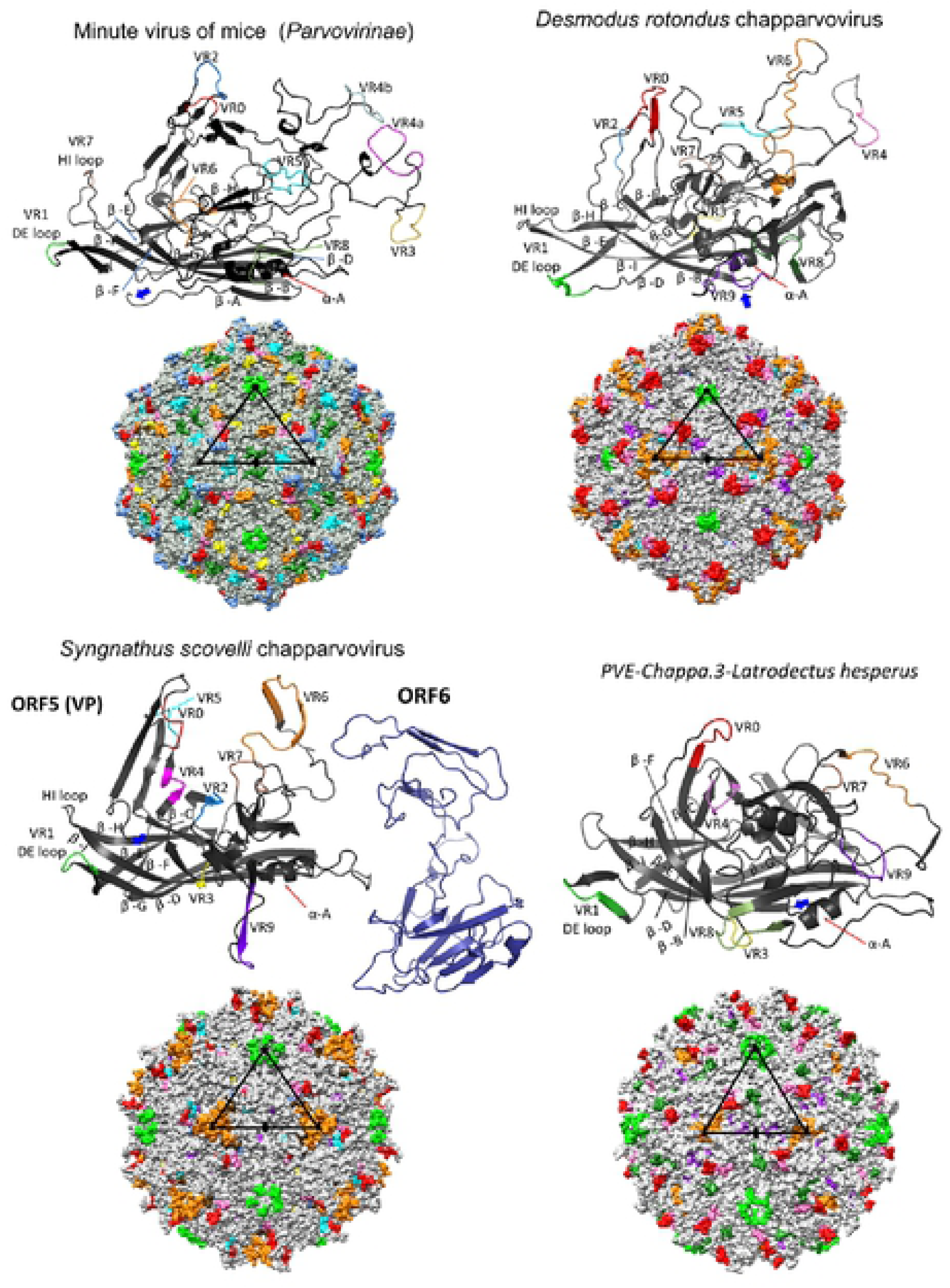
(A) Comparison of VP monomer ribbon diagrams of the protoparvovirus minute virus of mice (PDB ID: 1Z14) from subfamily *Parvovirinae* to homology models of an amniote, a fish and an endogenous arthropod chapparvovirus. Variable regions (VRs) of the same number are marked by the same color and mapped to the surface and luminal area of the icosahedral capsid model constructed of 60 monomers. In case of minute virus of mice, the VRs are marked by both the traditional numbering established for dependoparvoviruses (Latin numerals) and by the special one applied for protoparvoviruses only (Arabic numerals). Triangles mark the position of an asymmetric unit within the capsid, the fivefold symmetry axis marked by a pentagon, the threefolds with triangles and the twofold with an ellipsoid. (B) Homology model of ORF6, the hypothetical structural protein of *Syngnathus scovelli* chapparvovirus (ScChPV). The trimer of the ScChPV monomer model reveals a gap at each subunit interaction (arrows), unlike in case of the trimer of the hitherto smallest parvoviral capsid protein, *Penaeus stylirostris* densovirus. The gap might accommodate ORF6 in the assembled ScChPV capsid. Symmetry axes marked by the same symbols as for panel A.

In addition to their fundamental NS-NP-VP genome organization, chapparvoviruses encode various additional small ORFs. ORF1 is predicted to encode a small protein of approximately 15 kDa that contains a putative nucleus localization signal (NLS) in its C-terminal region. ORF1, which partially overlaps the N-terminal region of NS, is present in all genomes except PPV7. However, since the PPV7 genome also lacks the corresponding region of NS, this likely reflects a 5’ truncated genome sequence. The same is the case for turkey parvovirus (TPV2), although the C-terminal encoding region of the putative ORF1 protein could be revealed.

A second additional, putative ORF is present in only two of the chapparvoviruses examined here: PPV7 and simian parvo-like virus 3. This ORF, referred to as ORF2, is located downstream from ORF1 in a position completely overlapping the NS ORF. The TPV2 genome also contains a unique, presumably genome-specific additional ORF (ORF4) that overlaps the C-term encoding region of VP, and may encode a predicted 17 kDa protein (**Figure 2b**). Interestingly, this ORF was absent from the other, closely-related avian entries.

Analysis *in silico* revealed at least three potential promoters in chapparvovirus genomes. One of these is conserved throughout the clade, and is located upstream of all coding features, indicating that it likely drives early expression of virus genes. Moreover, its presence has been confirmed in MKDV by sequencing of cDNA derived from infected mouse tissue. None of the other potential promoters proved to be functional in case of MKDV, nevertheless. The MKDV transcriptome includes three confirmed transcripts to undergo splicing. Out of these, however, only the one with the shortest intron could be predicted to exist in case of all examined GenBank entries (**Figure 2b**). Interestingly, DrChPV, along with further rodent-derived ChPVs was predicted to possess the large intron of the putative VP transcript, hence displaying a strikingly MKDV-like transcription mechanism, despite of missing an acceptor site upstream the NP start codon.

In all ChPVs examined, with the exception of the 5’ truncated entries, we identified two potential polyadenylation signals in positions equivalent to those found in MKDV (Roedinger et al 2018), This implies that the polyadenylation strategy is a conserved feature of chapparvovirus transcription.

### Characterization of syngnathid chapparvoviruses and EPVs

We identified two chapparvoviral sequences in syngnathid fish. One of these, identified in the genome of the Gulf pipefish (*Syngnathus scovelli*) occurred within a relatively short scaffold (4002 nt). The entire scaffold was comprised of viral sequence, displaying truncated but nonetheless detectable J-shaped terminal hairpin-like structures (**Figure S1**). This suggests it likely represents a virus contaminant, as suspected for other ChPV sequences recovered from vertebrate WGS data (9). The virus from which this sequence was presumably derived was designated *Syngnathus scovelli* chapparvovirus (ScChPV). The ScChPV genome encodes a long NS ORF (807 aa), a strikingly short VP (367 aa) and a ChPV-like NP. Furthermore, a homologue of the ORF2 protein found in the amniote parvoviruses PPV7 and SPV3 was present. A predicted ORF was present in a genomic position equivalent to that of ORF1, found in amniote ChPVs. However, the predicted protein sequence did not disclose any detectable similarity to its amniote counterpart. ORF6, identified in partial overlap with the VP C-term encoding region, has the capability to encode a small protein of 27.2 kDa (239 aa), which demonstrated no sequential similarity to any GenBank entries to date. Fold recognition, however, revealed possible structural similarity to viral structural proteins, including the major envelope glycoprotein of Epstein-Barr virus (PDB ID: 2H6O chain A, p=0.012), the minor viral protein of the Sputnik virophage (PDB ID: 3J26, chain N, p=0.017) and the surface region of *Galleria mellonella* ambidensovirus (PDB ID: 1DNV, p=0.021) (**Figure 4**). These findings imply ORF6 may encode an additional structural protein in addition to VP.

A partial ChPV-like sequence identified in the genome of the tiger seahorse (*Hippocampus comes*) was flanked by extensive stretches of host genomic sequence, establishing that, unlike the ScChPV sequence identified in the Gulf pipefish genome assembly, it likely represents an EPV rather than a virus. Interestingly, however, phylogenies showed that both sequences obtained from syngnathe fish are relatively closely related, and cluster together with high bootstrap support (**Figure 3a and b**).

### Characterization of chapparvovirus-derived EPVs in invertebrate genomes

Via screening of WGS data derived from invertebrate species, we identified a total of 13 EPV sequences that disclosed a relatively close phylogenetic relationship to ChPVs. These elements showed varying degrees of degradation (**Figure 1).** In many cases only genome fragments were detected, and these usually included numerous nonsense mutations (**Table 1**). ChPV-derived elements were detected in three major arthropod clades that primarily occupy terrestrial habitats, namely arachnids of Chelicerata, chilopods of Myriapoda, as well as hexapod insects and entognaths.

Among the EPVs we identified, the most complete were identified in the Western black widow spider (*Latrodectus hesperus*). Two elements spanned the *rep, cap*, and NP genes, and a homologue of the *orf1* gene found in ScChPV (**Figure 2a**). At aa level of the putative NS1, however, these elements displayed only 62% identity. Although the *rep* of the element ChPV.2-Latrodectus hesperus appeared to encode the complete, 690-aa-long protein product, this ORF contained nonsense mutations in case of the other arachnid element, ChPV.3-Latrodectus hesperus (**Table 1**). In contrast, ChPV.3-Latrodectus hesperus displayed an undisrupted ORF1 of 113 aa, a slightly smaller homologue of its ScChPV counterpart as well as an intact cap, capable of encoding a 386-aa-long VP. In ChPV.2-Latrodectus hesperus only an N-term truncated ORF1 was present at the length of 37aa, along a *cap* disrupted by several nonsense mutations. Furthermore, the *cap* of this element was found to include an insertion of 74 aa, suspected to originate from a yet unknown repetitive element (revealed by sequence comparisons to be interspersed throughout the *L. hesperus* genome). ChPV.3-*Latrodectus hesperus* from the other hand, appeared to include an intact upstream region of the genome, revealing and additional small ORF of 81 aa length directly upstream the ScChPV ORF1 homologue, designated ORF1-Lh. This ORF disclosed no detectable homology to any entries to date. Upstream this ORF a potential promoter sequence could be revealed with high confidence (0.98 of 1). Both elements included complete, NP-encoding ORFs of 233 aa, although a canonical ATG start codon could be revealed only in case of one element (ChPV.3-*Latrodectus hesperus*).

We revealed the existence of two more elements in the Western black widow spider genome, although these only spanned the *rep* gene, disrupted by several nonsense mutations. However, one element (ChPV.4-*Latrodectus hesperus*) encoded nearly complete NS and NP genes, as well as a complete homologue of the ScChPV ORF1 gene. The true extent of preservation could not be assessed for this EPV as it occurred on a short scaffold that terminated within the EPV *rep* sequence. The putative NS1 was 0.8% identical to its counterpart in ChPV.2-*Latrodectus hesperus* at aa level. Interestingly, directly upstream this EPV another copy of the *rep* was present, encoding only the first 221 aa of the putative NS1 protein. This element also contained a complete ScChPV ORF1 homologue as well as ORF1-Lh, which did not display an ATG start codon in this case. The upstream promoter could only be identified with a much lower score of 0.6. ChPV.5-*Latrodectus hesperus* displayed a highly divergent, partial *rep* of 216 aa with only 0.42% identity to the ChPV.2-*Latrodectus hesperus* NS1 at aa level (**Figure 1**). This element clustered outside the monophyletic branch comprised by the three other EPVs of the same species (**Figure 3**).

A single chapparvovirus-derived EPV was identified in a second arachnid species - the Chinese golden scorpion (*Mesobuthus martensii*). This element was identified in a relatively short unplaced scaffold, and comparison to the *Latrodectus* elements indicated that the contig was truncated within the EPV, and consequently the true extent of its preservation could not be assessed. Nevertheless, ORFs disclosing homology to the NS, NP and VP proteins could be identified. While the first 100 or so codons of the NS ORF were absent, a complete NP ORF was detected, along with the first 46 codons of VP. All three ORFs were disrupted by frameshifts and stop codons (**Table 1**). No homologues of any alternative ORFs identified in other chapparvovirus genomes could be revealed.

We identified a chapparvovirus-derived EPV in the genome of a myriapod - the European centipede (*Strigamia maritima*). This element displayed partial homologues of the NS and NP encoding ORFs, both of which contained large deletions (**Figure 2b)** as well as numerous nonsense mutations (**Table 1**). Moreover, the NS ORF was disrupted by an extensive stretch of an insertion of unknown origin. No homologues of any of the above-mentioned alternative chapparvoviral ORFs could be identified in this endogenous sequence.

Seven chapparvovirus-derived EPVs were identified in hexapod arthropods (subphylum Hexapoda). One occurs in the genome of a bristletail species - the Northern forcepstail (*Catajapyx aquillonaris*) - belonging to the entognath order Diplura. The other six were identified in three species belonging to the vast insect order Coleoptera: the emerald ash borer (*Agrilus planipennis*), the taurus scarab (*Onthophagus taurus*), and the scarab beetle (*Oryctes borbonicus*). The bristletail element contains a C-terminal truncated *rep* of at least 250 aa and a near full-length NP ORF. The partial *rep* was intact, but the NP ORF is disrupted and highly divergent, showing significant sequence similarity only in the conserved core region of the putative protein. The ash borer element occurs in a scaffold that is ∼1 kb in length. One end of this scaffold contains 592 nt region exhibiting homology to the NS ORF, which harboured an N-terminal deletion of at least 200 aa.

In the case of the taurus scarab element ChPV.10-Onthophagus taurus, the almost complete NS ORF could be identified, disrupted by numerous nonsense mutations (**Table 1**). Interestingly, another endogenous element of parvoviral origin was detected at the same locus. This element encodes an intact, potentially fully-expressible NS gene, homologous to the NS1 of ambidensoviruses (genus *Ambidensovirus*) and discloses similarity to a recently reported ambidensovirus species, detected only at cDNA level in the transcriptome of two bumble bee species (*Bombus cryptarum* and *B. terrestris*) (22). An additional ORF was present in this ambidensoviral element, overlapping the reputative NS1 gene, which harboured no significant similarity to any sequences deposited to GenBank to date. In its derived aa sequence, however, a homeobox domain could be revealed, also in intact, potentially expressible state. The other two elements of the taurus scarab genome located in the same assembly scaffold, only 2540 nts apart from each other. Both EPV consisted of only a partial ORF, which disclosed similarity to chapparvoviral *rep*s. None of these elements encompassed the tripartite helicase domain, hence they were not included in phylogenetic inference.

Two EPVs were derived from the scarab beetle genome. One of these, designated ChPV.13-*Oryctes borbonicus*, harboured a near complete *rep* at 402 aa as well as a short, partial *cap*, capable of encoding only the first 33 aa of the putative VP. The region of rep homology occurred within an ORF that was not disrupted by any frame shifts and extended without disruption upstream and downstream, suggesting a putative longer gene product - potentially encoding a longer, divergent NS protein - to be present. However, these regions did not disclose sequence similarity to any proteins hitherto deposited to GenBank. The *Oryctes* element (EPV-ChPV.14-Oryctes), in contrast, included only a heavily truncated NS of 254 aa (**Figure 1**).

### Structural characteristics of chapparvovirus capsids

We built 3D homology models to facilitate comparison of chapparvovirus capsid structures to those found in other parvoviruses. The derived polypeptide sequence of the complete VP ORF encoded by DrChPV was subjected to fold recognition, to identify suitable templates for homology modelling. This comparative analysis showed that the most similar VP protein occurs in parvovirus H1 (genus *Protoparvovirus*) (PDB ID: 4G0R, p=9e-05), and this sequence was therefore used as the template for homology modelling. To overcome the stochastic aspect of model construction, due to the lack of sequence identity and the non-homologous nature of the ChPV VP genes to other parvoviral VPs, we used the final model obtained in this analysis as a template for analysis of four further ChPVs - rat parvovirus 2, PPV7, TPV2, and pit viper chapparvovirus. The pitfalls of using models as templates has to be noted here, however, this method ensured that only those regions showed structural variability, which would probably do so in case of the actual capsid structures.

We examined the VP sequences of two representatives of the second major ChPV clade (**Figure 3b**) – one derived from a presumably exogenous virus (ScChPV) and one from an EPV (ChPV.3-*Latrodectus*). Fold recognition identified, for the VP protein encoded by ScChPV dependoparvovirus VP proteins as potential templates (adeno-associated virus 8, PDB ID: 2QA0, p=8e-04; Adeno-associated virus rh32.33, PDB ID: 4IOV, p=9e-04), while for the VP encoded by the black widow spider EPV the most reliable hit was the VP4 protein of an iteradensovirus (*Bombyx mori* iteradensovirus, PDB ID: 3P0S, p=8e-04). When superimposing the obtained models with the VPs of AAV8 and BmDV, however, structural similarity only covered the jelly roll core and the αA helix, whereas only the EC loop out of the traditionally more variable surface loops.

Modelling indicated that the ChPV VP monomer harbours an eight-stranded β-barrel ‘jelly roll’ core and the αA helix at the twofold symmetry axis as found in all members of the family *Parvoviridae* to date (23) (**Figure 4a**). Equivalents of all short strands were present (β-C, H, E, F), as well to four out of the five longer strands (β-B, D, I, G). However, no structural analogue to the outmost β-A could be identified (**Figure 4**). Examining the secondary structure prediction results further disproved the existence of a β-A analogue, indicating β-B to be the closest to the N-term. The first strand of the *Syngnathus scovelli* ChPV VP, appeared to fold outside of the jelly roll, leaving the longer side of the barrel without a β-B, comprised of only three strands, namely D,I and G, despite of a complete upper, CHEF sheet (**Figure 4**). When modelling the complete T=1 capsid polymer, this manifested as a hole, which is normally covered by β-B even in case of the smallest parvoviral capsids (**Figure 4b**). All ChPVs as well as the only ChPVe VPs displayed two canonical loops surrounding their fivefold axes, linking sheets D with E at the channel and sheets H with I on the floor surrounding the channel. In case of the amniote ChPVs, the pore displayed a tight opening. The sequence of the DE loop varied to some extent among these seven entries, which also manifested in the models. The HI loop was, however, highly conserved throughout, containing only one variable position between the amniote ChPVs.

We mapped the chapparvoviral VRs identified by VP alignments (**Figure S3**) to both VP monomers and complete capsids, to examine how they manifest on the virion surface and make comparisons to parvoviruses of known structure, represented by the minute virus of mice (MVM) capsid structure, as the prototype virus of subfamily *Parvovirinae* (PDB ID: 1Z14) (**Figure 4**). Out of ten chapparvoviral VRs identified (VR 1 to 10), presented by **Figure S3a** only VR1, VR2, and VR9 proved to be similarly positioned, hence probably analogous to the parvoviral counterparts. Some VRs appeared to be positioned at luminal surface of the chapparvovirus capsid (VR4 in all cases and VR8 of the amniote ChPVs, whereas VR6 of the *Latrodectus* element and ScChPV), unlike all parvoviruses studied to date with the exception of bovine parvovirus, a bocaparvovirus (24) in which VRVIII shows this configuration. Since the ChPV VRs appeared to be non-homologous to those established for either proto- or dependoparvoviruses, we re-defined them by numbering from N to C-term.

In addition to their distinctive VRs, ChPVs ubiquitously appeared to harbor a highly variable C-terminal region, with a length varied between 12 to 62 residues. The ChPV VP variable C-term appears to be buried in most cases with the exception of ScChPV, where it is probably exposed. In case of the Latrodectus element, it forms the luminal surface of the threefold, whereas in the case of fish and amniote ChPVs it is located at the twofold (**Figure 4a**).

The *ScChPV* and ChPV.3-*Latrodectus hesperus* VP lacked a VR6 homologous to that of the amniote ChPVs albeit displayed variation in another position instead, still in the sixth-place counting from the N-term (**Figure S3b**). Moreover, both of them displayed truncated VRs 3, 5, and 7, compared to their amniote counterparts. VR9, furthermore, was absent from the ScChPV VP, whereas VR 10 was missing from the VP of ChPV.2-*Latrodectus hesperus* (**Figure S3b**). As for the surface, the largest variable region for amniote ChPVs, namely DrChPV is VR7, forming the entire three-fold protrusions, with VRs 1, and 9, forming small protrusions surrounding the aforementioned peaks.

The complete capsid models of non-amniote chapparvoviruses were observed to harbor surface features that are strikingly distinct from those of the amniote ones, more closely resembling the capsids of the *Ambidensovirus*-*Iteradensovirus* clade of *Densovirinae* (see **Figure 3a**), with a surface that is less spikey (**Figure 5**).

**Figure 5.**
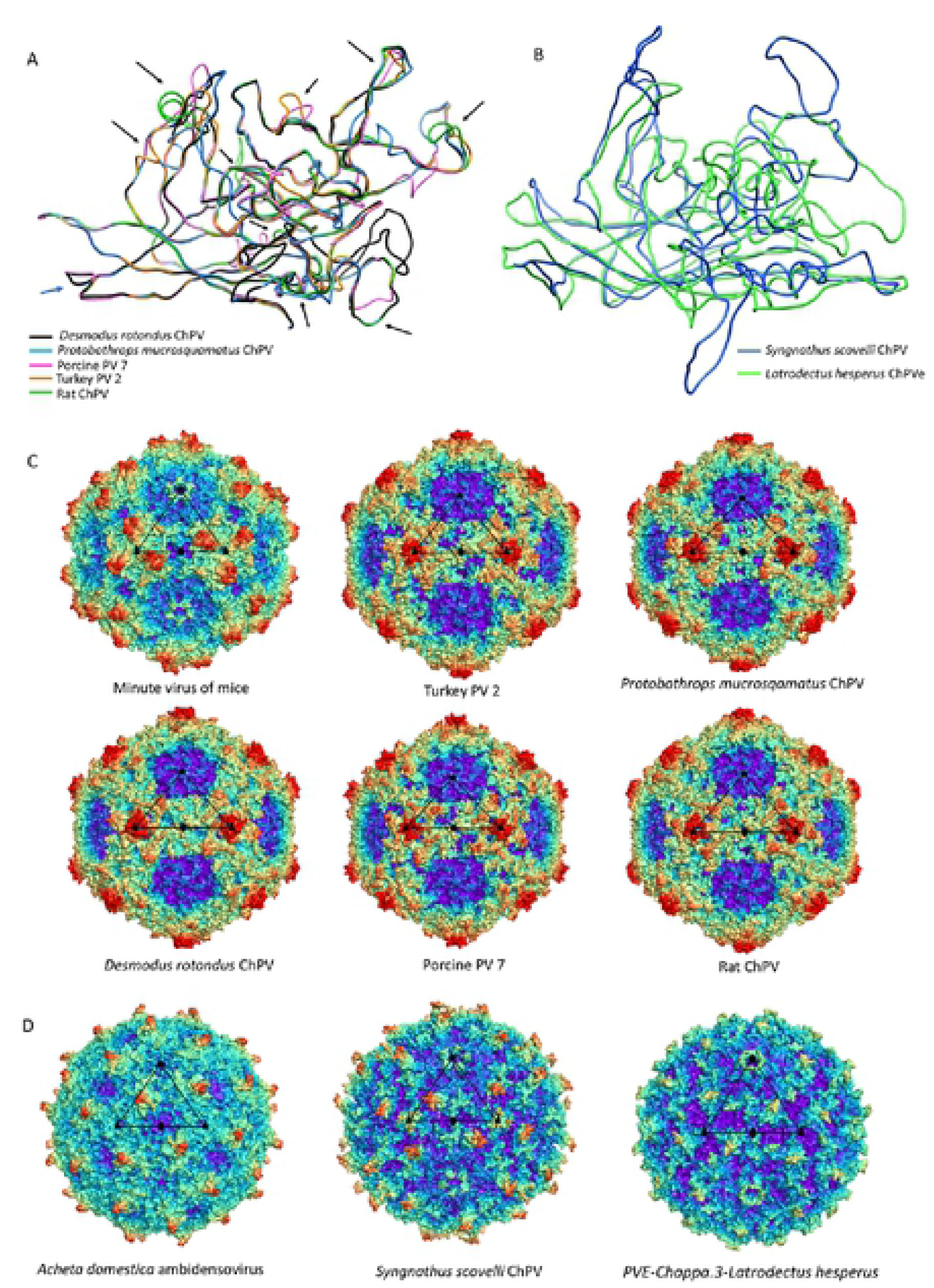
(A) The three different predicted chapparvovirus VP structural types represented by ribbon diagrams. The first panel shows sSuperposition of protein monomer homology models of amniote chapparvovirus capsids, including reptilian, avian, rodent, chiropteran and ungulate representatives. Black arrows show variable regions (VRs) The next two panels show the homology model of a fish chapparvovirus capsid monomer and an arthropod endogenous chapparvoviral one. (B) Capsid surface morphology of amniote chapparvovirus homology models compared to that obtained for a prototypic parvovirus, minute virus of mice (MVM) (PDB ID: 1Z14 at 3.25 Å resolution). Capsids are orientated by their twofold symmetry axes, as shown in the line diagram. Below the comparison of homology models of complete viral capsid surface morphology of the newly-identified fish chapparvovirus and arachnid endogenous chapparvoviral element is shown with that of the actual capsid structure of two densoviruses (subfamily *Densovirinae*, genus *Ambidensovirus*) (PDB ID: 4MGU at 3.5 Å resolution for *Acheta domestica* ambidensovirus and 1DNV at 3.7 Å for Galleria mellonella ambidensovirus).

We also constructed a homology model of ORF6 of ScCHPV, based on the minor viral protein of the Sputnik virophage (PDB ID: 3J26). This protein appears to harbor multiple beta strands close to its C-term, out of which the outermost could potentially fill in the aforementioned gap, caused by the lack of a βB (**Figure 4b**).

## DISCUSSION

Historically, the family *Parvoviridae* has always been comprised of two subfamilies, with specificity for vertebrate or invertebrate hosts being the major demarcation criterion (2). This division was initially supported by phylogenetic inference, however, as the number of densoviral genera increased, the heterogeneity of densoviruses, specifically their segregation into two clades, has not gone unnoticed (1). Our study provides further evidence that the traditional division of parvoviruses into vertebrate-specific and invertebrate-specific subfamilies no longer holds - rather, it supports the division of the *Parvoviridae* into four major subgroups: the *Parvovirinae*, a split *Densovirinae*, and the Chapparvoviruses as illustrated in **Figure 3a**.

The data presented here show that chapparvoviruses infect an exceptionally broad range of hosts, including both vertebrates and invertebrates. Furthermore, we show that chapparvoviruses found in fish are more closely related to those that infected ancestral arachnoid arthropods than they are to those that infect amniote vertebrates such as rodents (**Figure 3b**). These findings suggest that chapparvoviruses have been transmitted between distantly related host species in the past.

Phylogenies indicate that all amniote ChPVs have a common origin (**Figure 3a**), consistent with the overall conservation of their genome organization and some aspects of predicted transcriptional strategy (**Figure 2b**). Previous studies suggested that these viruses might have broadly co-diverged with host species (9). The present, expanded data set implies that some transmission of ChPVs between distantly related vertebrate classes may have occurred among (**Figure 3b**) - though it should be kept in mind that almost all amniote ChPVs have been identified via metagenomic sequencing of environmental samples, mostly fecal viromes, thus their true host affiliations remain uncertain.

Sequences derived from parvoviruses occur relatively frequently in animal genomes (13, 14, 17, 25, 26). However, these sequences overwhelmingly derive from a small proportion of parvovirus lineages. For example, ambidensovirus-derived EPVs dominate invertebrate genomes (14), whereas vertebrate EPVs almost exclusively derive from dependoparvoviruses, protoparvoviruses, or viruses closely related to these two genera (12, 13, 25, 26). In this study we found no trace of ChPV-derived sequences in tetrapod genomes, despite recent evidence that ChPVs infect this host group (21). By contrast, ChPV-derived sequences are relatively common in arthropods, with the genomes of some species harboring multiple, independently acquired ChPVs (**Table 1**). The tendency of EPVs to derive from a subset of parvovirus genera likely has biological underpinnings. For example, in vertebrates it may reflect the tendency of dependoparvoviruses to integrate into host DNA, and/or the requirement of protoparvoviruses to initiate DNA damage response (DDR) during replication (27, 28). Similar features of the viral life cycle could account for the biased distribution of ChPV-related sequences in species genomes - i.e. arthropod and fish ChPVs might have adopted a replication strategy that favors germline integration, whereas that of amniote ChPVs precludes it. Notably, some arthropod species have integration sites containing multiple independently acquired EPVs of both ChPV and ambidensovirus origin, suggesting that hotspots of parvovirus integration and/or fixation might exist in their genomes.

Our discovery of ChPV-derived elements in fish and arthropod genomes establishes that ChPVs can infect these species in addition to mammals (21). Moreover, it provides evidence that the ChPVs are probably an ancient lineage of parvoviruses. Although we did not identify any orthologous ChPV insertions, the EPVs described here show extensive evidence of germline degradation. Through comparison to studies of EPVs in mammals (in which several orthologous EPVs have been described), it appears likely that chapparvoviruses have been present in animals for many millions of years. Moreover, as the hexapod EPVs appear to be monophyletic and mirror the evolution of their host species, the age of ChPVs could possibly correlate with the Insecta-Entognatha split, suggesting a minimum age of 400 million years (29).

Through comparative analysis of EPVs and chapparvoviruses, we show that chapparvovirus genomes exhibit a number of defining characteristics. Firstly, all possess a short, monosense genome, encoding a relatively large NS and a relatively short VP. The short VP proteins of ChPVs are clearly homologous to one another, but show no similarity to those found in other parvovirus lineages. Similar to those found in the penstyl-, hepan- and brevidensoviruses, the VP proteins of ChPVs lack PLA2 domains. Notably these are also the genera to which ChPVs are most closely related in NS-based phylogenies (**Figure 3a**).

Secondly, chapparvoviruses typically encode multiple additional gene products besides the replicase and capsid. To begin with, almost all encode a nucleoprotein (NP) gene in an overlapping frame with *rep*. In this report, we show that putative NP ORFs are present in ChPV-derived EVEs, suggesting it is an ancestral, conserved feature of these viruses. It’s absence, however, from the coleopteran lineage is intriguing, as it is still present in the EPV of the hexapod stem group Diplura of Entognatha. Phylogenetic reconstructions (and the extensive overlap with *rep*) imply it was acquired ancestrally and independently lost in the lineage derived from members of the hexapod crown group, Coleoptera (**Figure 3b**).

ChPV genomes appear to encode several predicted auxiliary ORFs in addition to NP. A functional role for auxiliary ORF1 is supported by (i) it’s conservation across the entire amniote ChPV clade, and; (ii) limited experimental data indicating it is expressed in MKPV via a spliced transcript. Auxiliary ORF2 was only identified in a small subset of ChPV genomes, but a functional role for this ORF is suggested by the presence of homologues in distantly related ChPVs of amniotes and fish (see **Figure 3b**). Interestingly, although all ChPVs appear to express ORF1 via splicing of a small intronic sequence (**Figure 2**), those harboring an ORF2 homologue are predicted to lack the peculiar large introns found in expression of MKPV NP and VP transcripts (21). ScChPV lacks an ORF1 homolog, but contains a predicted reading frame in the corresponding position. Homologues of this ScChPV ORF1 variant are present in all three arachnid EPVs, although not in the first, but in the second position. As only the three *Latrodectus* EPVs possess a homologue of ORF1-Lh, it is possible that this small ORF became incorporated only after the split from the syngnathe fish lineage, whereas the incorporation of ScChPV ORF1 predates it. The distribution of homologous auxiliary genes across phylogenetic lineages of ChPVs implies that distinct lineages have acquired and/or lost these genes on multiple, independent occasions.

MKPV has been reported to possess only one promoter and two polyadenylation signals, as well as an extensive number of spliced transcripts. This transcription pattern, however, appears to be unique to only one lineage of the amniote ChPVs, comprising of rodent, chiropteran, primate, and reptilian entries. As both the avian ChPV sister clade as well as the lineage including PPV7 appear to display genome organization specific for these groups suggesting that ChPVs infecting host groups outside of the MKPV lineage may utilize distinct transcription strategies.

Despite the potential pitfalls of homology modeling, and the use of distinct templates to reconstruct both the VP monomer and capsid structures, we obtained remarkably similar predicted structures for VP sequences found in closely related viruses/EPVs. Since the viral capsid plays an important role in mediating the interactions between parvoviruses and their hosts, comparisons of capsid structures can potentially reveal insights into parvovirus biology. Homology modelling indicated that ChPV VPs would assemble into a complete, T=1 icosahedral capsid, despite their relatively small size. Furthermore, their predicted structures are remarkably similar to that found in other parvoviruses, despite the lack of any detectable similarity in the sequences of their VP proteins. Similarities include the presence of a conserved jelly roll core and αA helix, the existence of the DE and HI loops, and the presence of identifiable VRs. Interestingly, the amniote ChPV capsids appear to possess the same number of VRs as most of the vertebrate parvoviruses of subfamily *Parvovirinae*, even if only a few of them (namely VRs 1, 2 and 9) proved to be analogous features. In these virus capsids, variations were most prominent among the threefold peaks and protrusions as well as the two-fold depression, as observed in members of the *Parvovirinae* (**Figure 5**). The tendency of some VRs to manifest at the luminal surface of the capsid in models suggests these regions could play a role in intracellular host-virus interactions. However, in order to become accessible to interact elements of intracellular signaling pathways, either uncoating would be necessary, or potential conformational changes to expose these regions. Based on previous findings, however, the parvovirus capsid appears to traffic into the nucleus intact (30, 31). Considering this, these buried regions might play a role in processes linked to the nucleus. Interestingly, however, bovine parvovirus, the only other parvovirus in which buried VRs have previously been observed (32), is an enteric pathogen, and metagenomic studies suggest that amniote ChPVs (with the exception of MKPV) are also enteric.

In addition to the VRs, all ChPVs seem to harbor highly-variable VP C-terms. A similar phenomenon has been observed in case of iteradensoviruses, where the last 40 C-terminal residues are disordered and their structure could not be resolved (33). Although the location of the ChPV C-term appears to vary, its association with regions that are overtly involved in parvovirus-host interactions (e.g. the two- and threefold peaks) is certainly intriguing.

MKPV is associated with pathology of the urogenital system, whereas a related virus - murine ChPV - has been detected at very high prevalence in murine liver tissue suggesting it is a gastrointestinal agent (34). The VPs of the two, however, only differ in six aa residues, located within VR 3 and near VR2 on the surface and in the buried VR4 as well as in the similarly buried variable C-term (**Figure S3a**). Thus, these positions could constitute potential determinants of tissue tropism in murine ChPVs. Parvoviruses subfamilies *Parvovirinae* and *Densovirinae* utilize distinct strategies to stabilize their icosahedral capsids (35). Vertebrate parvoviruses extend the longer side of the jellyroll fold with an additional, N-terminal strand by folding back β-A to interact with the twofold axis of the very same monomer, hence creating an extended ABDIG sheet (36, 37). By contrast, the densovirus capsid preserves the symmetric arrangement of the jellyroll fold, and possesses a β-A which is a direct elongated N-terminal extension of the β-B instead, interacting with the β-B strand of the neighboring monomer toward the fivefold axis (38, 39). Strikingly, our data show that ChPV capsids lack β-A strands (and also the β- B strand in the case of ScChPV). The functional implications of this are unclear - possibly ChPV capsids are stabilized in the absence of β-A via a yet unknown, additional VP. If ChPVs express additional structural proteins, they are presumably encoded by spliced transcripts (given the unusually small size of the *cap* gene). Alternatively, the ChPV capsid might assemble without the incorporation of an additional β strand, perhaps at the cost of losing the stability and resilience typical of parvoviruses in general. Potentially, this could account for the apparent presence of buried VRs. Interestingly, in studies of MKPV, viral proteins could be detected in the kidneys of infected mice even though no assembled particles could be observed in inclusion body-affected tubular cells (21). This, along with our structural predictions, suggests that the ChPV strategy for uncoating and cellular trafficking might be very different from that found in the *Parvovirinae* and *Densovirinae*.

Uniquely, the genome of ScChPV appears to include a putative additional structural protein (ORF6), in addition to the above-mentioned alternative ORFs. All parvoviruses to date - except those of genus *Penstyldensovirus* (39) - have been reported to incorporate up to three additional minor capsid proteins into the virion, which share a common C- terminal region. To encode a structural protein on an entirely separate ORF sharing no mutual coding sequence with *cap* would be unique. Possibly, this unusual feature could be connected to the predicted lack of a β-B strand in ScChPV VP protein.

Taken together, the data presented here establish that the ChPVs belong to a parvovirus lineage that comprises a distinct lineage from all other parvoviruses, and infects an exceptionally broad range of host species, including both vertebrates and invertebrates. Consistent with this, their relatively complex genomes exhibit numerous unique features, implying their life cycle might significantly differ from what has been established in case of other members of the family. These findings underscore the need for further basic and comparative studies of ChPVs, both to assess their potential impact on animal health, both wildlife and livestock. Furthermore, this is the first study to imply that vertebrate parvoviruses are not monophyetic, as well as members of the family must have evolved to infect vertebrates on at least two separate occasions.

## METHODS

### Genome screening and sequence analysis

WGS data were screened for chapparvovirus sequences using the database-integrated genome screening (DIGS) tool (40). ChPV sequences were characterized and annotated using Artemis Genome Browser (41). The BLAST program was used to compare sequences and investigate predicted viral ORFs. To determine potential homology and sequence similarity even between previously undescribed ORFs, we constructed a local database by including all ORFs exceeding 100 aa in length derived from all the exogenous and endogenous sequences incorporated into this study and used the local BLAST P and X algorithms to conduct similarity searches in it. Two ORFs were accepted as homologous if they gave a significant hit in case of an expectation value threshold of 1.

Promoters were predicted using the neural network-based promoter prediction server of the Berkeley Drosophila Genome Project and further verified by the Promoter Prediction 2.0 server (42, 43). Splice sites were also detected using the neural network-based applications of the Berkeley Drosophila Genome project and SplicePort (43, 44). Polyadenylation signals were predicted by the SoftBerry application POLYAH (45). To verify the suitability of these applications to be capable of detecting the above mentioned chapparvoviral transcription elements we ran MKDV through the workflow pipeline.

### Phylogenetic reconstructions

The derived aa sequences of ORFs disclosing homology to parvoviral NS1 proteins were aligned with at least five representatives of each genera of Parvoviridae, or with one representative of each species of given genus in case the number of species did not exceed five. To ensure the correct identification of the tripartite helicase domain, structural data was also incorporated into alignment construction using T-coffee Expresso (46) and Muscle (47). The full-length NS1 derived aa sequences of the chapparvovirus clade were aligned by Muscle and the M-coffee algorithm of T-coffee (48). Model selection was carried out by ProTest and the substitution models RtREV+I+G in case of the helicase-based inference and LG+I+G for the complete chapparvoviral NS1 tree were predicted to be the most suitable based on both Akaike and Bayes information criteria. To infer the maximum likelihood phylogenetic tree the PhyML-3.1 program was used with 100 bootstrap iterations (49), based on a guide tree previously constructed by the ProtDist and Fitch programs of the Phylip 3.697 package (50).

### Homology modelling and DNA structure prediction

Structural homology was detected by applying the pGenTHREADER and pDomTHREADER algorithms of the PSIPRED Protein Sequence Analysis Workbench (51). The same workbench was used to map disordered regions using DISOPRED3 and to predict the secondary structure of the complete chapparvoviral VP protein sequences via the PSIPRED algorithm. The selected PDB structures were applied as templates for homology modeling, carried out by the I-TASSER Standalone Package v.5.1 (52). To guide the modelling, the predicted secondary structures were applied as a restriction. 60-mers of the acquired purative VP monomer structures were constructed by the Oligomer Generator feature of the Viper web database (http://viperdb.scripps.edu/) (53). Surface images of the capsids were rendered using the PyMOL Molecular Graphics System (54). Capsid surface maps and VP monomer superposition were carried out by UCSF Chimera (55). To predict the presence of potential DNA secondary structural elements the DNA Folding Form algorithm of the mFold web server was utilized (56).

## Acknowledgements

RJG was funded by the Medical Research Council of the United Kingdom (MC_UU_12014/12). WMS is supported by the Fundação de Amparo à Pesquisa do Estado de São Paulo, Brazil (Scholarships No. 17/13981-0). MAM and JP are funded by NIH R01 GM109524.

## Supporting Information Legends

**Figure S1**

Secondary structure predictions of the *Syngnathus scovelli* chapparvovirus genome termini.

**Figure S2**

(A) Secondary structure and disordered region predictions of the nucleoprotein (NP) ORF from an amniote chapparvovirus (mouse kidney virus) and an endogenous chapparvoviral element from an invertebrate genome (ChPV.3-Latrodectus hesperus). (B) Predictions of potential phosphorylation sites in case of the same amniote chapparvovirus and an invertebrate endogenous chapparvoviral element NPs.

**Figure S3**

(A) Alignment of amniote chapparvovirus capsid protein ORF (VP) derived amino acid sequences, containing both isolates of murine origin. Variable regions (VRs) are marked by the black bars and coloring is based on sequence similarity (red = highly similar, blue = not similar). The conserved loops making up the fivefold symmetry axes of the capsid are highlighted in bold. (B) The same alignment incorporating the complete VP protein sequences of all novel chapparvovirus sequences reported in this study.

